# Coronary Artery Disease risk variant dampens the expression of CALCRL by reducing HSF binding to shear stress responsive enhancer in endothelial cells

**DOI:** 10.1101/2023.02.08.527795

**Authors:** Ilakya Selvarajan, Jin Li, Petri Pölönen, Tiit Örd, Kadri Õunap, Aarthi Ravindran, Kiira Mäklin, Anu Toropainen, Lindsey K. Stolze, Stephen White, Casey E. Romanoski, Merja Heinäniemi, Yun Fang, Minna Kaikkonen-Määttä

**Affiliations:** A. I. Virtanen Institute for Molecular Sciences, University of Eastern Finland, P.O. Box 1627, 70211 Kuopio, Finland; Institute of Biomedicine, School of Medicine, University of Eastern Finland, P.O. Box 1627, FIN-70211, Kuopio, Finland; Department of Cellular and Molecular Medicine, The College of Medicine, The University of Arizona; Tucson, AZ 85721, USA; Department of Medicine, The University of Chicago, Chicago, IL 60637, USA; Department of Life Sciences, Manchester metropolitan university, Manchester, M1 5GD, UK

**Keywords:** Coronary artery disease, GWAS, SNP, HAEC, endothelial, shear stress, HSF, CRISPR, STARR-Seq, RNA-Seq

## Abstract

Coronary artery disease (CAD) is one of the major causes of mortality worldwide. Recent genome-wide association studies have started to unravel the genetic architecture of the disease. Such efforts have identified Calcitonin receptor-like (CALCRL), an important mediator of the endothelial fluid shear stress response, associated with CAD risk variants. In this study we functionally characterized the non-coding regulatory elements carrying CAD risks SNPs and studied their role in the regulation of *CALCRL* expression in endothelial cells. We demonstrate that rs880890-harboring regulatory element exhibits high enhancer activity and significant allelic bias with A allele showing 40% more activity than G allele. We also observed that the A allele of rs880890 is favored over the G allele under shear stress. CRISPR deletion of rs880890-enhancer resulted in downregulation of *CALCRL* expression. EMSA further showed that heat shock factors are binding to the enhancer with a preference for A allele over the G allele. In line with this, HSF1 knockdown resulted in a significant decrease in *CALCRL* expression. *CALCRL* knockdown as well as variant perturbation experiments confirmed the role of CALCRL in the regulation of eNOS, apelin, angiopoietin, prostaglandins and endothelin-1 signaling pathways while demonstrating a significant decrease in cell proliferation and tube formation. Overall, our results demonstrate the existence of an endothelial-specific heat shock factor regulated transcriptional enhancer carrying a CAD risk SNP rs880890 that regulates *CALCRL* expression. Better understanding of *CALCRL* gene regulation and the role of SNPs in modulation of *CALCRL* expression could provide important steps towards understanding genetic regulation of shear stress signaling responses.

## Introduction

Coronary artery disease (CAD) is a disease of coronary vessels that develops due to build-up of atherosclerotic plaques in the vessel wall. Genome-wide association studies (GWAS) have identified ~300 risk loci for CAD, that are beginning to shed light into the complexity of its genetic architecture^1–5^. Although some associations locate within coding regions, approximately 98% of signals come from non-coding regions, which suggests that single nucleotide polymorphisms (SNPs) within gene regulatory elements play major role in mediating the effects on gene expression^6–8^.

One of the identified CAD GWAS loci on chromosome 2q32.1 harbors the calcitonin receptor-like (*CALCRL*) gene^1^, encoding for a seven-transmembrane G-protein-coupled receptor that mediates the pleiotropic effects of calcitonin gene-related peptide (CGRP) and adrenomedullin (ADM)^9^. These two structurally related neuropeptides were originally described as potent vasodilators. Beyond blood pressure regulation, CALCRL is involved in a variety of key biological processes, including cell proliferation, modulation of apoptosis, vascular biology, and inflammation, and is currently emerging as a novel target for the therapy of migraine^10^. In solid tumors, antibody-mediated inhibition of CALCRL signaling has been demonstrated to reduce tumor growth via disruption of angiogenesis or via direct antiproliferative effects on cancer cells^11^. Interestingly, it was further shown that CALCRL is expressed in normal CD34+ hematopoietic progenitors and that CGRP and ADM directly act on CD34+ cells to promote colony formation *in vitro*, indicating a functional role of CALCRL in physiological myelopoiesis^12^

It was recently demonstrated that *CALCRL* is a major mediator of nitric oxide (NO) pathway in endothelial cells through endothelial fluid shear stress response^13^. A fundamental function of the endothelial layer of blood vessels is their ability to sense fluid shear stress and to transform this information into intracellular signals^14^. These mechano-sensing and -signaling processes are critical for maintaining vascular integrity, thus affecting physiological vascular tone and morphogenesis but also pathological vascular processes including hypertension and atherosclerosis. In the present study, we demonstrate that the non-coding regulatory region in the 3’ end of *CALCRL* is important for gene expression and show that rs880890, variant associated with both hypertension and CAD risk, exhibits allele specific regulatory activity. We further demonstrate that stress regulates this region through binding of Heat Shock Factor (HSF) transcription factors and silencing of HSF1 significantly reduces CALCRL expression. We also confirm that the “A” allele of the SNP increases HSF1 binding over the “G” allele, leading to an allelic bias. Overall, our study suggests a causal role for rs880890 in disrupting the vasodilator, angiogenesis and eNOS pathway in which CALCRL plays a key mediator role.

## Material and method

### Culture of cell lines

TeloHAEC (ATCC, CRL-4052), HEPG2 (ATCC, HB-8065), RAW 264.7 (ATCC TIB-71), pre-adipocyte 3T3-L1 (ATCC CL-173) and MOVAS (ATCC CRL-2797) cell lines were used in the luciferase experiment. TeloHAEC were cultured using Vascular Cell Basal Medium (ATCC PCS-100-030) supplemented with Endothelial Cell Growth Kit-VEGF (ATCC PCS-100-041) and 10 % FBS. HepG2 cells were cultured in Dulbecco’s modified Eagle medium (DMEM; 4.5 g/L glucose, 2 mM L-glutamine, LONZA) supplemented with 10% fetal bovine serum (FBS; GIBCO). RAW 264.7 were cultured using DMEM (DMEM), and FBS: (FBS in water bath at 56 degree for 30 minutes). 3T3-L1 was cultured using DMEM (Lonza BE12-614F) supplemented with 10% FBS and 2mM L-glutamine. MOVAS was cultured using Dulbecco’s Modified Eagle’s Medium (DMEM) supplemented with geneticin. Primary human coronary artery endothelial cells (HCAECs) were cultured in media MV2 (purchased from Promocell) and used between passage 4-6. 100 U/ml penicillin, 100 μg/ml streptomycin was used in all cell-lines used in this study to prevent bacterial contamination.

### Gene and eRNA expression quantification

Gene transcript coordinates were identified from GRO-seq data using a custom de novo detection pipeline^15,16^. eRNA quantification was performed using Homer^17^ analyzeRepeats.pl, with parameters: -noadj -noCondensing -pc 3. Quantification was done from the opposite strand of coding gene transcription for intragenic enhancers and from both strands for intergenic enhancers. Gene end coordinates were expanded by 1500bp to account for transcription at the end of transcripts and annotating intragenic gene enhancers. Counts for eRNA and gene expression were normalized using limma voom^18^. Only enhancers with cpm > 1 in at least 5 samples were kept.

### Elastic net regression

Elastic net multivariable regression model was fitted for each gene to identify enhancers that are the best predictors of gene expression. For each gene, only enhancers that were in the same TAD and P-value < 0.05 were used for fitting the model. Alpha parameter, controlling the L1+L2 regularization proportion was set to 0 to 1 with 0.05 increments. Caret^19^ and glmnet^20^ R packages and repeatedcv method with 5 repeats and folds set to 5 were used for fitting and cross validating the model. Model with best fit after tuning the alpha and lambda parameters was kept. Enhancers were ranked based on their importance for the model, as determined by the model coefficient.

### CRISPR-Cas9–Mediated Deletion of Enhancer in TeloHAECs

The CRISPR reagents were adapted from the Alt-R system from IDT. The guide RNAs were designed using the IDT-tool to minimize off-targeting effects using two guides to create 372-bp deletion (**Table S1**). The positive control (crRNA targeting HPRT) from the Alt-R CRISPR-Cas9 Human Control Kit (IDT) were used as control. Delivery of the ribonucleoprotein complex into TeloHAEC was performed using Neon transfection system with a 1350 V pulse of 30 ms width. Forty-eight hours post transfection RNA was purified using a RNeasy Mini Kit (Qiagen) and the cDNA was prepared with a RevertAid First Strand cDNA Synthesis Kit (Thermo Fisher Scientific). The mRNA level of the CALCRL was measured by SYBR green chemistry qPCR using specific primers (**Table S2**) in a StepOne real-time PCR system (Thermo Fisher Scientific). Three independent experiments with 4 technical replicates were performed. Data (ΔCt values) was checked for normal distribution before performing statistical tests. Paired Student’s T-test (two-tailed) was used for data that followed normal distribution and equal variance. Otherwise, Mann–Whitney U test was used. p < 0.05 was used to define a significant difference between the groups.

### Single-cell RNAseq data processing and analysis

scRNA-Seq data previously generated from atherosclerotic human coronary arteries^21^ was obtained from the NCBI GEO repository (accession GSE131778). The data processing and cell type annotation was carried out as described in^22^ using Seurat package^23^. For plotting gene expression in individual cells, the gene counts were depth-normalized to 10,000 total counts per cell and log-transformed.

### Dual Luciferase reporter assays

DNA fragments containing the region rs880890-rs840585 with haplotypes (G-T) and (A-C) were amplified by PCR from genomic DNA (gDNA) with Phusion polymerase and specific primers **(Table S3)**. For the dual luciferase reporter assay, amplified DNA (698bp) was subcloned into hSTARR-seq_ORI plasmid (Addgene, #99296) that was used as a backbone for the plasmid constructs. The enhancer inserted plasmid was co-transfected into TeloHAEC, HEPG2, RAW 264.7, pre-adipocyte 3T3-L1 and MOVAS cell lines with the control vector pGL4.75 (Promega), which encodes the luciferase gene hRluc (Renilla reniformis). The luciferase constructs prepared with the inserts were verified by sequencing. Transfection was performed in 6-well plate using Lipofectamine Stem Transfection Reagent (Thermofisher STEM00008) according to the protocol. Three independent experiments with four technical replicates were performed. Paired Student’s T-test (two-tailed) was used for data that followed normal distribution and equal variance. Otherwise, Mann–Whitney U test was used.

### STARR-Seq allelic reporter assay processing

TeloHAEC and HepG2 cell lines were used for the experiment. STARR-Seq library and experiment have been described previously in^22,24,25^. To assess transcriptional activity of haplotypes of the candidate regulatory regions, STARR-Seq RNA read data was UMI-deduplicated, depth-normalized to library size, and normalized for haplotype abundance in the plasmid DNA (input) library.

### Motif analysis of rs880890

To identify the potential transcription factors binding to the region with rs880890, we used PERFECTOS-APE^26^ using TFBS motif collection using P-value cut-off of 0.0005 and fold change cutoff of 2.0 and motifbreakR^27^ package with default settings.

### Endothelial cells ChIP-Seq/ATAC-Seq data analysis related to shear stress

Since TeloHAECs are homozygous to the region of the SNP rs880890, the following ChIP-Seq data related to endothelial cells (HUVEC) cultured under shear stress (for 6 h at 37°C and with constant CO2 perfusion at ± 3 dyn/cm2) and static was downloaded from GSE116241. Sequencing data was downloaded from NCBI using the SRA Toolkit and processed using the nf-core ChIP-seq pipeline^28^ to derive bam files. The BaalChIP^29^ tool was used for allele-specific measurements of transcription factor binding from the ChIP-Seq data. A Bayesian statistical approach was used by the tool to correct the effect of background allele frequency on the observed ChIP-Seq read counts. ATAC-Seq data related to Human aortic endothelial cells (HAEC) that were heterozygous at rs880890 undergoing shear stress were downloaded from GSE112340. Reads were aligned to hg19 with Bowtie2^30^. SAMtools^31^ was used to convert sam to bam files. BAM files were sorted and indexed using SAMtools sort and index function. IGV viewer^32^ was used to visualize the SNP position and count up the read counts in unidirectional (UF) and disturbed flow (DF).

### HSF1 knockdown and CALCRL expression under shear stress

Human aortic endothelial cells (HAECs) were seeded in 6-well plates at 100% confluency and cultured in flow media with EGM-2 medium (Lonza) containing 4% dextran. HAECs were subjected to unidirectional flow (UF) or disturbed flow (DF) for 24 hours^33^ before being harvested for RNA isolation. HAECs were seeded in 6-well plates in EGM-2 medium at 90% confluency and were transfected with 50 nM siRNA using RNAiMAX (Life Technologies) as described by the manufacturer. siRNA targeting HSF1 (SI03246348) and non-targeting control (1027310) were both purchased from Qiagen. Media was changed the following day to flow media with EGM-2 medium containing 4% dextran. HAECs were subjected to unidirectional flow (UF) for another 24 hours^33^ before being harvested for RNA isolation. RNA was isolated from cells using GenEluteTM Mammalian Total RNA Miniprep Kit (Sigma Aldrich) and reverse transcribed using High-Capacity cDNA Reverse Transcription Kit (ThermoFisher). Quantitative mRNA expression was determined by RT-qPCR using SYBR Green MasterMix (Roche). The mRNA level of the HSF1 knockdown under shear stress was measured by using specific primers (**Table S4**).

### TeloHAEC nuclear extraction

TeloHAEC cells were collected (5 × 10^6^) in PBS by scraping from culture flasks and washed twice with cold PBS. The re-suspension of the cells was 500 μl 1x Hypotonic Buffer (20 mM Tris-HCl, pH 7.4, 10 mM NaCl and 3 mM MgCl2) with protease inhibitor cocktail by pipetting up and down several times and incubated on ice for 15 minutes. 25 μl detergent (10% NP40) was added and vortexed for 10 seconds at highest setting. Centrifugation was carried out of the homogenate for 10 minutes at 3,000 rpm at 4°C was carried out. The nuclear pellet was re-suspended in 50 μl complete Cell Extraction Buffer (10 mM Tris, pH 7.4, 2 mM Na3VO4, 100 mM NaCl, 1% Triton X-100, 1 mM EDTA, 10% glycerol, 1 mM EGTA, 0.1% SDS, 1 mM NaF, 0.5% deoxycholate and 20 mM Na4P2O7) for 30 minutes on ice by vortexing at 10 minute intervals followed by sonication and centrifugation for 30 minutes at 14,000 x g at 4°C. Quantitation of protein concentration was performed using the Quant-iT™ Protein Quantitation Kit (Thermo Fisher Scientific, Rockford, USA), the aliquoted supernatant (nuclear fraction) was stored in -80°C.

### EMSA

Oligonucleotide probes (**Table S5**) (15 bp flanking SNP site for reference or alternate allele) with a biotin tag at the 5′ end of the sequence (Integrated DNA Technologies) were incubated with TeloHAEC nuclear extract and the working reagent from the LightShift™ Chemiluminescent EMSA kit (Thermo Fisher Scientific, Catalog number:20148, Rockford, USA). For competitor assays, an unlabeled probe of the same sequence was added to the reaction mixture at 100 × excess. The reaction was incubated for 30 min at room temperature, and then loaded on a 6% retardation gel (Invitrogen, Catalog number: EC6365BOX, California). The contents of the gel were transferred to a nylon membrane and visualized by UV trans illumination Image Lab Software (Bio-rad). For supershift experiments, 1.5ul of HSF1 antibody (SBT) were added before addition of the oligos.

### siRNA mediated knockdown of CALCRL

For siRNA treatment, TeloHAEC were reverse transfected with Lipofectamine RNAiMAX transfection reagent (Thermo Fisher) according to the manufacturer’s instructions. The target-specific Silencer Select siRNAs for *CALCRL* and scrambled siRNA were obtained from Thermo Fisher (s19889 and s19890). Cells were lysed 24 h after transfection. Three independent experiments with 4 technical replicates were performed. RNA extraction, cDNA synthesis, qPCR and statistical analysis followed the protocol described above for CRISPR-Cas9 experiments.

### RNA-Sequencing and Data Analysis

For the *CALCRL* silenced and enhancer deleted samples, total RNA quality was assessed using the Agilent 2100 Bioanalyzer System. RNA library preparation was handled using the QuantSeq 3′ mRNA- Seq Library Prep Kit FWD (Lexogen, Vienna, Austria) according to the manufacturer’s instructions. The library was quantified using Qubit dsDNA HS Assay Kit (Q32854, ThermoFisher Scientific) and its quality was checked with Bioanalyzer using High Sensitivity DNA Kit (5067-4626, Agilent Technologies). Individual libraries were pooled in equimolar ratio (4 nM for each) and sequenced on an Illumina NextSeq 500. The bcl2fastq2 v2.20 (Illumina) was used to demultiplex sequencing data and to convert base call (BCL) files into FASTQ files. Nf-core^28^ RNA-seq pipeline was used to process fastq files and derive gene counts. Gene counts were normalized for effective library size, and differentially expressed genes (DEGs) were analyzed using DESeq2^34^. DEGs were defined by a P value of <0.05 and an absolute fold change of >2.

### Expression of downstream target genes under shear stress

HCAECs were seeded on 0.1% gelatin coated slides with a density of 2.5 × 10^5^ and cultured for a minimum of 3 days to ensure complete confluency, production and reorganisation of sub-cellular matrix and maturation of cell-cell junctions. Culture under flow was performed for 72 hours using a parallel plate flow apparatus as described^35–37^ under disturbed flow (±0.5Pa, 1 Hz OSS), or unidirectional flow (1.5Pa LSS) shear stress. Total RNA was extracted from HCAECs using the RNeasy Mini Kit (Norgen) according to the manufacturer’s protocol. Total RNA was extracted from HCAECs using the RNeasy Mini Kit (Norgen) according to the manufacturer’s protocol. 100ng RNA were reverse transcribed into cDNA with random primer by reverse transcriptase (RT) (Qiagen). cDNA (relative 1.67ng RNA) was amplified by standard qPCR with Taq DNA polymerase (Sensifast, SybrGreen, LOW-ROX Kit, Bioline). The geometric mean of GAPDH, RPLP0 and PABC4 quantification was used to normalise the expression of each test gene. Paired Student’s T-test (two-tailed) was used for data that followed normal distribution and equal variance. Otherwise, Mann–Whitney U test was used. p < 0.05 was used to define a significant difference between the groups (n=7 individual donors).

### Network analysis and identification of CALCRL co-expressed genes

The WGCNA^38^ (Weighted gene correlation network analysis) was used for identification of co-expressed genes from RNA-Seq samples. WGCNA is a well-established tool for studying biological networks based on pair wise correlation between variables in high-dimensional RNA-Seq data sets. ComBat-seq^39^ was used to remove batch effect from the data. Sequencing data was processed like mentioned in the data analysis section. After rlog transformation, we identified the genes and samples with excessive numbers of missing samples were identified using the function goodSamplesGenesMS and were discarded from the analysis. The samples were then clustered using the function hclust to identify outliers in the dataset. We constructed a weighted gene network using soft thresholding power β. The function pickSoftThreshold was used that aids in choosing a proper soft-thresholding power. We chose a power of 10, which resulted in an approximate scale-free topology network with the scale-free fitting index R2 greater than 0.9. For the remaining genes, the choice of the soft thresholding power β was used to which co-expression similarity is raised to calculate adjacency with function adjacency(). The matrix of correlations was converted to an adjacency matrix of connection strengths. To minimize effects of noise and spurious associations, the adjacency was transformed into Topological Overlap Matrix using TOM() and the corresponding dissimilarity was calculated. The function hclust() was used to hierarchically cluster the genes. Branches of the dendrogram group together densely interconnected are highly co-expressed genes. Modules were then defined as sets of genes with high topological overlap. The module hub gene was defined as the gene in the module with the highest connectivity or based on a high intra-modular connectivity. Cytoscape^40^ app geneMANIA^41^ was used to identify known and novel co-expressed genes.

To identify how much of the co-expressed genes are enriched in the differential expressed dataset of *CALCRL* depletion, a gene enrichment analysis was performed using the Hypergeometric t-test^42^. The parameters for the hypergeometric t-test are as follows. Number of successes (k=49) was the co-expressed genes significantly differentially expressed under *CALCRL* depletion (P-value<0.05), sample size (s=195) was the total co-expressed genes, number of successes in the population (M= 1460) represented all the genes that were significantly differently expressed under *CALCRL* depletion (P-value<0.05; in both siRNA and enhancer deletion) while population size (N= 11820) was the total number of genes included in the analysis.

### Xcelligence for studying proliferation

Transfected cells were seeded at 5,000 cells per well in 200 µl of complete media in E-plates (ACEA Biosciences, San Diego, CA, USA) and grown for 48 hours while monitored with an xCELLigence DP system (ACEA Biosciences) which measures electrical impedance across interdigitated gold micro-electrodes integrated on the bottom of tissue culture plates. The xCELLigence recorded cell index readings every 1 hour for 3 days. Three biological replicates were used to determine cell proliferation and represented the relative numbers of cells compared to control cells. A two-way ANOVA test was used to compare *CALCRL* siRNA knockout treatment to scrambled control; with P ≤ 0.05 deemed significant.

### Tube Formation Assay

Forty-eight–well plate was coated with Matrigel basement membrane matrix 150 μL/well (Corning, growth factor reduced). For CALCRL knockdown, TeloHAECs were seeded on to the Matrigel 48 hours after siRNA transfection. Knockdown efficiencies of more than 90% were consistently attained. Images were taken from 9 spots per replicate every 4 hours until 24 hours using Incucyte (S3 System). Each siRNA knockdown had 3 technical replicates in each experiment, and experiments were repeated 3 times. Analysis was done with ImageJ^43^ and associated macro tool Angiogenesis Analyzer^44^ where total tube length was selected as the parameter of evaluation. For time-course analysis, each time point represents the average tube length from all spots of all replicates with same treatments at the same time point. Statistical analyses were performed with two tailed t test.

## Result

### Identification of *CALCRL* regulatory enhancer by co-expression analysis

CALCRL is a G protein-coupled receptor and a recently identified CAD locus^11^. Further analysis of the SNPs within the *CALCRL* locus using Open Target Genetics-portal^45^ demonstrated that the SNPs are also associated with hypertension (**Figure 1A**). Since majority of the variants were located within non-coding regions and overlapped enhancer elements marked by DNase hypersensitivity and H3K27ac, we next sought to identify the relative importance of individual enhancers in the regulation of *CALCRL* expression. Starting from the premise that enhancer (e)RNAs are co-expressed with their target genes^46,47^, we constructed co-expression networks based on regularized logistic regression. This was achieved by comparing co-activity patterns of eRNAs and coding gene expression across cell types based on public GRO-Seq data on 336 samples representing 45 cell types^15^. Our model reports a linear combination of coefficients derived from eRNA expression to describe how important each enhancer is as predictor for gene expression. Overall, twelve regulatory regions (**Figure 1B, C**) were included in our analysis and enhancer-8 demonstrated the highest regression coefficient of 0.31 and was thus predicted to be central for the regulation of *CALCRL* expression **(Table S6)**. Hi-C data from TeloHAECs^48^ further confirmed the interaction between enhancer-8 and the promoter of *CALCRL* (**Suppl. Figure 1**). Finally, to confirm the predicted regulatory role of enhancer-8 in controlling endothelial *CALCRL* expression, the Alt-R CRISPR-Cas9 system was used to selectively delete a 372-bp genomic region in TeloHAECs using electroporation. PCR assays detected around 50% genetic deletion in regulatory region enclosing rs880890 in TeloHAECs (**Supp. Figure 2**). The enhancer deleted cells exhibited significantly lower gene expression (P=7.5E-5) compared to control cells confirming the importance of the enhancer elements in regulating *CALCRL* expression (**Figure 1D**).

**Figure 1:**
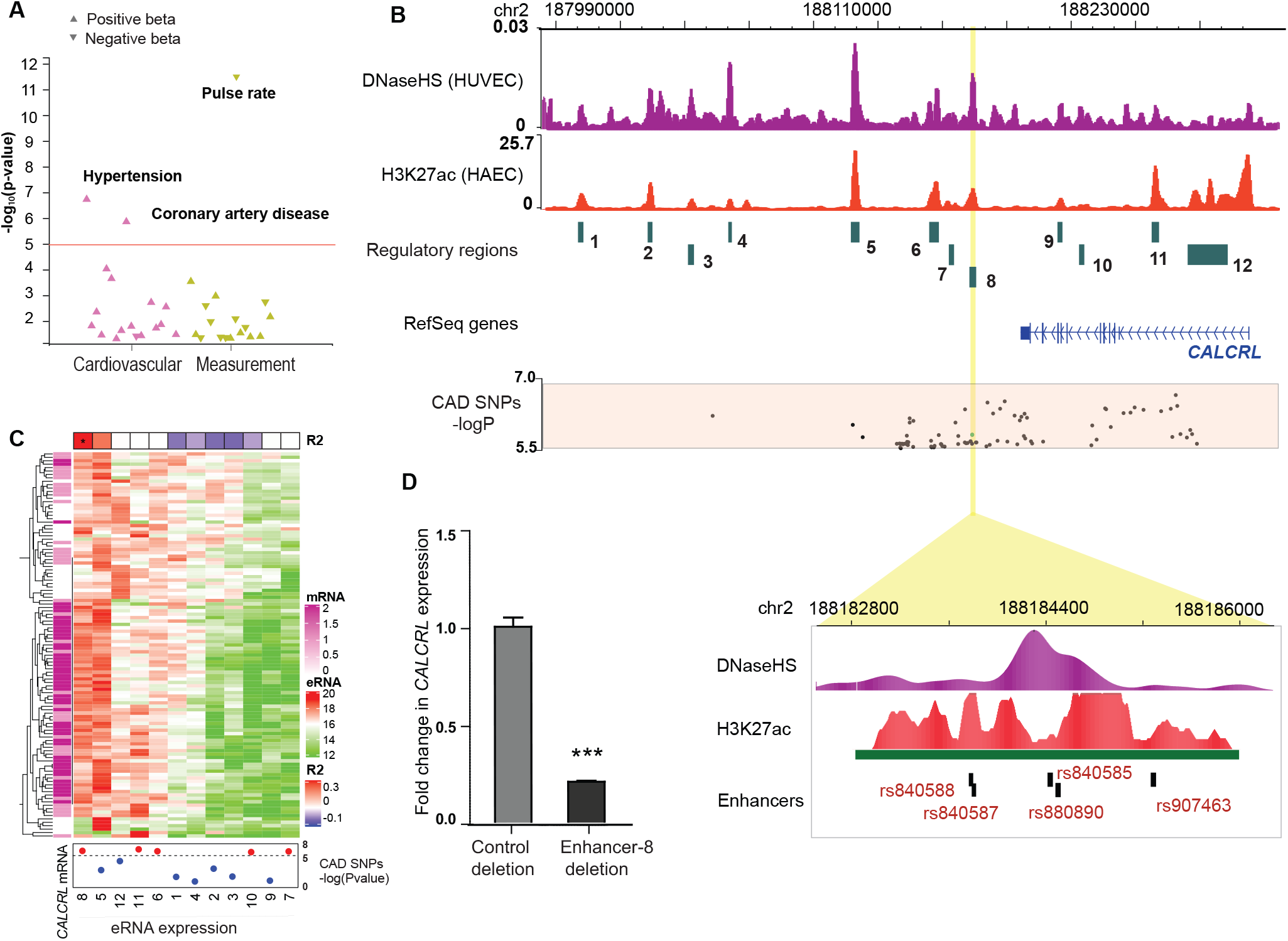
**A)** Association of rs880890 with CVD related traits from Open Targets Genetics-portal44. rs880890 was significantly associated with hypertension and CAD. **B)** WashU browser shot of CAD loci with *CALCRL* region showing the regulatory regions (1-12) included in the study. A close-up look of the enhancer-8 harboring CAD associated SNPs. DNAse hypersensitivity shows that region with rs880890/rs840585 is accessible compared to nearby regions **C)** Heatmap showing the eRNA expression of each enhancer included in the study, *CALCRL* mRNA expression, R2 value of enhancers and the CAD SNPs annotations. **D)** Results of CRISPR-Cas9–mediated regulatory region deletion in TeloHAECs. qPCR was performed from three independent experiments. The statistical significance was evaluated using a two-tailed Student’s t-test or Mann–Whitney U test. For the bar plot, significance is denoted with an asterisk. *** p<0.0005.

**Figure 2:**
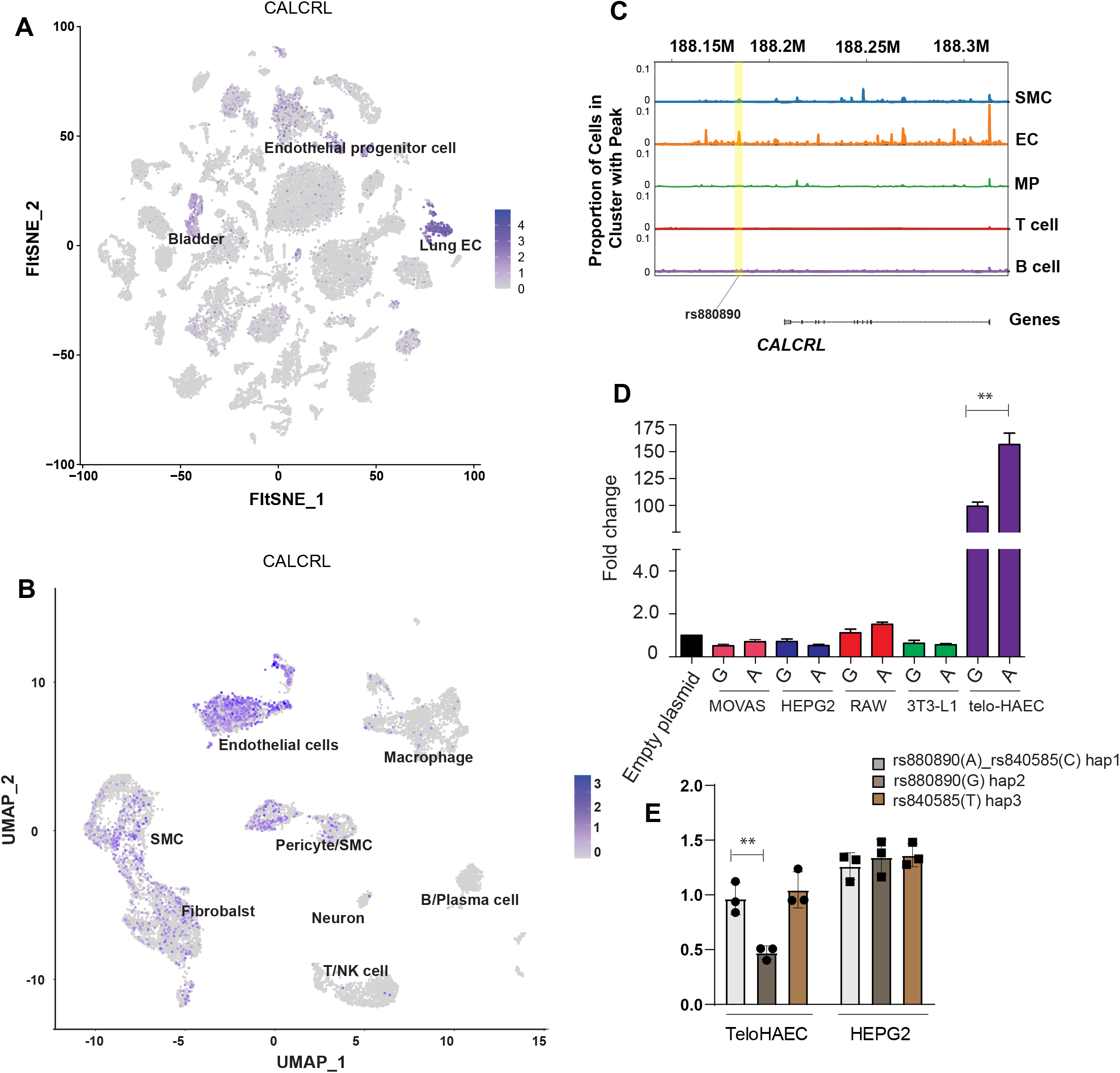
**A)** Single cell RNA-Seq data from Tabula muris representing data accross 20 organs and tissues from mice shows that *CALCRL* is highly expressed in endothelial cells, especially in the lung. **B)** Single cell RNA-Seq data from human coronary arteries^21^ demonstrating that CALCRL is predominantly expressed in endothelial cells followed by smooth muscle cells and fibroblasts. **C)** Pseudobulk coverage track visualization of single nucleus ATAC-Seq signal^22^ at the *CALCRL* loci showing endothelial cell type-specific activity of the rs880890 containing regulatory element. **D)** The activity of the rs880890-containing enhancer was investigated in MOVAS, HEPG2, RAW 264.7, 3T3-L1 and TeloHAECs. Luciferase assay showing a significant increase in enhancer activity in region carrying rs880890 “A” allele compared to “G” allele in TeloHAEC followed by RAW 264.7 while other cell types did not show enhancer activity. **E)** STARR-Seq results from TeloHAEC and HepG2 showing the enhancer activity in the presence of rs880890 or rs840585. Results show that difference in enhancer activity was observed only in the presence of rs880890 in TeloHAEC and no difference in enhancer activity in HepG2 was seen (n=3).

### Allele specific activity of rs880890-harboured enhancer in Telo-HAEC

The enhancer-8 harbored five CAD associated SNPs, and we next focused on characterizing rs880890 and rs840585 in the DNAse hypersensitivity region in more detail. **(Figure 1B zoom out)**. GTEX database suggested that *CALCRL* expression is ubiquitous throughout tissues **(Suppl. Figure 3)** with the highest expression seen in lung, artery, and adipose tissue. To identify the cell type contributing to the expression of *CALCRL* in the arterial tissue, we took advantage of single cell data from mouse single cell atlas (*Tabula muris*^49^). tSNE visualization of all tissues from *tabula muris* identified *CALCRL* expression predominantly in endothelial cells (**Figure 2A**). In addition, analysis of public scRNA-Seq data from human atherosclerotic lesions^21^ demonstrated high expression of *CALCRL* in endothelial cells and to lesser degree in smooth muscle cells (**Figure 2B; Supplementary Figure 4**). In line with the cell type specific gene expression, enhancer-8 was only found accessible in endothelial cells of the human atherosclerotic aorta based on our previously published single nucleus ATAC-Seq data^22^ (**Figure 2C**).

**Figure 3.**
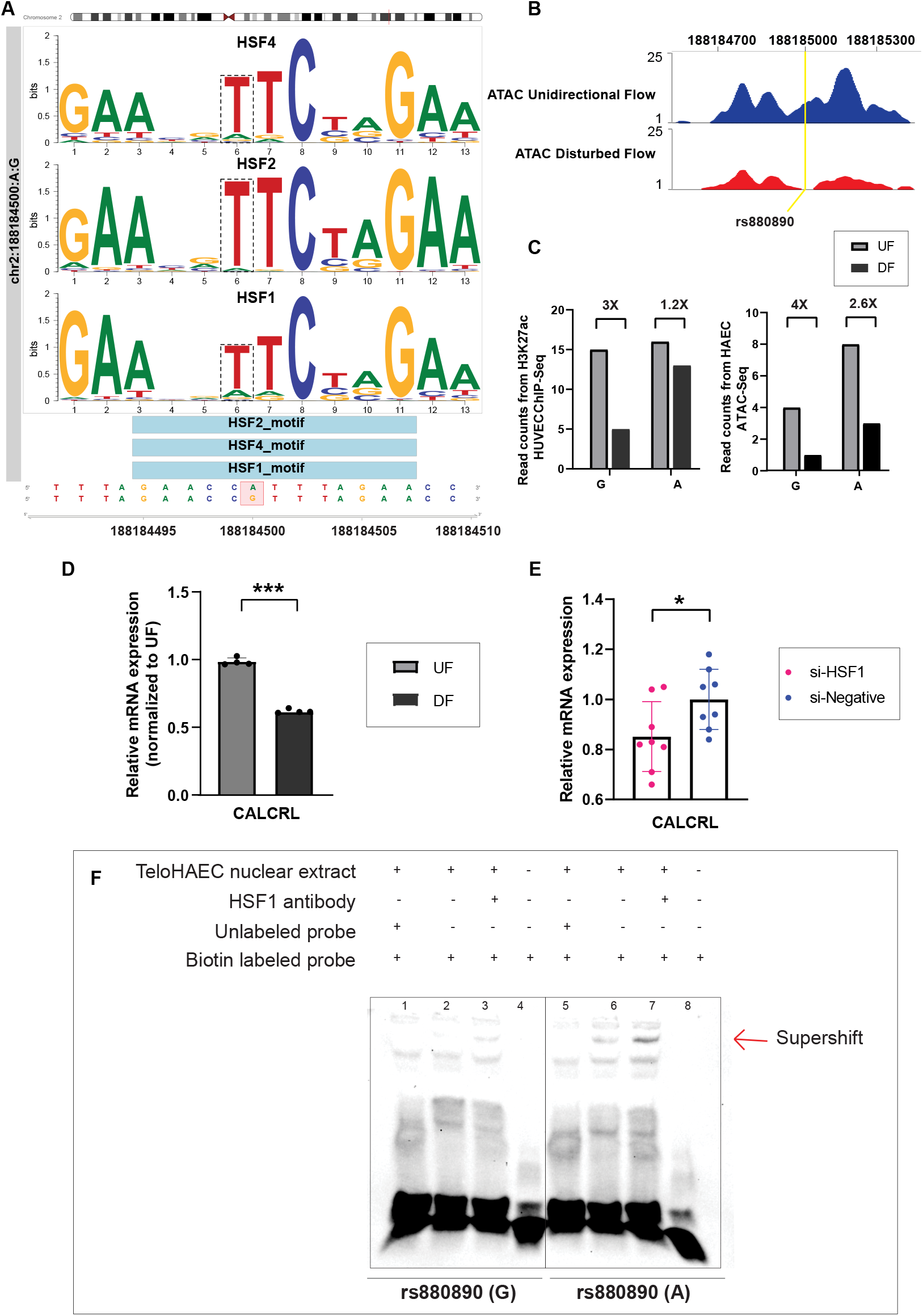
**A)** Motif analysis showing the binding affinity of rs880890 ‘A’ and ‘T’ allele compared to ‘G’ allele. **B)** ATAC-seq demonstrating higher accessibility of the rs880890 cis-regulatory element (enhancer) in HAECs under unidirectional flow (UF) compared to disturbed flow (DF). **C)** Increased ATAC-seq and ChIP-Seq reads in rs880890-containing region in HAECs under UF compared to DF. Bar plot shows increased ATAC/ChIP-Seq reads in cells under UF and lower reads from the G allele-containing chromosome under DF compared to A allele. **D)** A significant decrease (P<0.0001) in the expression of *CALCRL* was detected under DF compared to UF. **E)** Knockdown of HSF1 under shear stress resulted in significant downregulation of *CALCRL* expression (P=0.02) **F)** EMSA showing a supershift for HSF1 indicative of increased binding in the presence of “A” allele. The statistical significance was evaluated using a two-tailed Student’s t-test or Mann–Whitney U test. For the bar plot, significance is denoted with an asterisk. *P < 0.05; *** p<0.0001.

To study if the cell type specificity could be driven by the enhancer-8, we measured the enhancer activity of 698bp region in endothelial cells (TeloHAEC), pre-adipocytes (3T3L1), smooth muscle cells (MOVAS), hepatocytes (HepG2) and macrophages (RAW264.7). Enhancer-8 harbored two CAD SNPs in the DNase hypersensitive region, and both were cloned into the luciferase plasmid as a haplotype (Ref: rs880890(A)_rs840585(C) and Alt: rs880890(G)_rs840585(T)). Low enhancer activity was detected in all cell types except TeloHAECs which demonstrated 100-fold higher activity compared to the empty plasmid. Importantly, when we compared the luciferase signal of “A” allele compared to the “G” allele, there was a 40% increase in the activity of “A” compared to “G” **(Figure 2D)**. This *cis*-eQTL was confirmed in GTEx database where rs880890 “A” allele was significantly associated with increased expression of CALCRL compared to “G” allele. Also supporting our results, a trend of allele specific deposition of histone mark H3K27ac and the binding of endothelial specific transcription factor ERG was also detected in a cohort of 21-44 HAEC donors^50^ **(Suppl. figure 5)**.

To separate between the two SNPs in mediating the allele specific effect, we cloned the naturally occurring three haplotype combinations into a reporter vector with haplotype 1 containing the reference alleles rs880890(A) and rs840585(C) while haplotype 2 carrying rs880890(G) allele and haplotype 3 the rs840585(T) allele. Our analysis suggests that compared to haplotype 1, only haplotype 2 showed allele specific enhancer activity while hap3 showed no effect **(Figure 2E)** pinpointing rs880890 as the most probable causal SNP. Importantly, rs880890 also demonstrated significant evolutionary conservation (PhyloP -log p-value = 0.57) compared to rs840585 that was predicted to be fast evolving (PhyloP -log p-value = -0.99).^51^

### Heat shock factors bind to CALCRL regulating enhancer

Sequence based motif analysis revealed that heat shock factors such as HSF1, HSF2 and HSF4 are likely to bind to the *CALCRL* regulating enhancer at the rs880890 SNP region with a preference towards the “A” allele in binding affinity **[Figure 3A; Table S7]**. HSFs are known to activate heat shock proteins during stress^52^ and in vascular endothelial cells, HSF1 increase corresponds to elevated levels of eNOS^53^. Since CALCRL is a key mediator of eNOS pathway implicated in fluid shear stress response, and HSF1^54^ has been shown to be activated by shear stress, we wanted to investigate the effect of rs880890 SNP on enhancer activity under such conditions. ATAC-seq analysis of HAECs subjected to unidirectional (athero-protective) flow similar to hemodynamics in human distal carotid artery or the disturbed flow (DF) waveform which resembles the hemodynamics in human carotid demonstrated that the rs880890-harboring enhancer is more accessible in HAEC under UF compared to DF (**Figure 3B**). Moreover, analysis of the publicly available H3K27ac ChIP-Seq data allowed us to interrogate the allelic bias of rs880890 under shear stress in endothelial cells. The results confirmed that “A” allele is indeed biased towards higher enhancer activity binding compared to “G” allele and this effect is more prominent under shear stress in HAECs and HUVECs **(Figure 3C; Table S8)**. These effects were concordant with the regulation of *CALCRL* expression which was higher in HAECs subjected to UF and lower upon DF (P<0.001) (**Figure 3D)**. Additionally, HSF1 knockdown under unidirectional flow (Figure 3E) resulted in a significant decrease in *CALCRL* expression. This is in line with the directionality of HSF1 expression under shear stress with lower HSF1 levels detected under DF compared to UF^55^.

To confirm the predicted allele specific binding of HSF1 to the rs880890, we performed electromobility shift assay (EMSA) using biotinylated 31-bp probes targeting either the reference or alternative allele in TeloHAECs. Competitor assays were performed by incubating the reaction with ×100 excess of unlabeled (no biotin) oligonucleotide complexes with identical sequence. The results demonstrated an increased protein binding to the “A” allele when compared to the “G” allele of rs880890. To show that the shift in band in due to HSF binding, we further performed a supershift experiment with HSF1 antibody. Indeed, a supershift was detected and increased binding of HSF1 binding to the “A” allele was recorded. **(Figure 3G)**. Together these results suggest that the rs880890 SNP harboring enhancer is indeed responsive to shear stress, and this is likely mediated by the allele specific binding of HSF1.

## CALCRL knockdown and enhancer deleted cells affects vasodilation and endothelial specific pathways

To understand the downstream effects of *CALCRL* and *CALCRL* regulating enhancer, we performed RNA-Seq on upon siRNA-mediated *CALCRL* knock down and CRISPR-mediated rs880890-enhancer deletion. We then used the rlog transformed matrix of our RNA-Seq data to construct co-expression network using 11820 genes across 22 samples. This allowed identification of 31 distinct gene co-expression modules in our RNA-Seq data depicted in distinct colors (**Suppl. figure 6**). The *CALCRL* gene was found in the module blue along with 143 co-expressed genes. Dendrogram of consensus module eigengenes obtained on the consensus correlation was used to merge the similar modules blue and cyan with a threshold of 0.1 resulting in identification of 195 genes co-expressed with *CALCRL*. Selected genes are illustrated on a geneMANIA network (**Figure 4A**) that demonstrated the relationships between CALCRL and co-expressed genes. Interestingly, we observed potentially new *CALCRL* co-expressed genes such as *APLN, ADAMTS18* and *AKAP12*.

**Figure 4:**
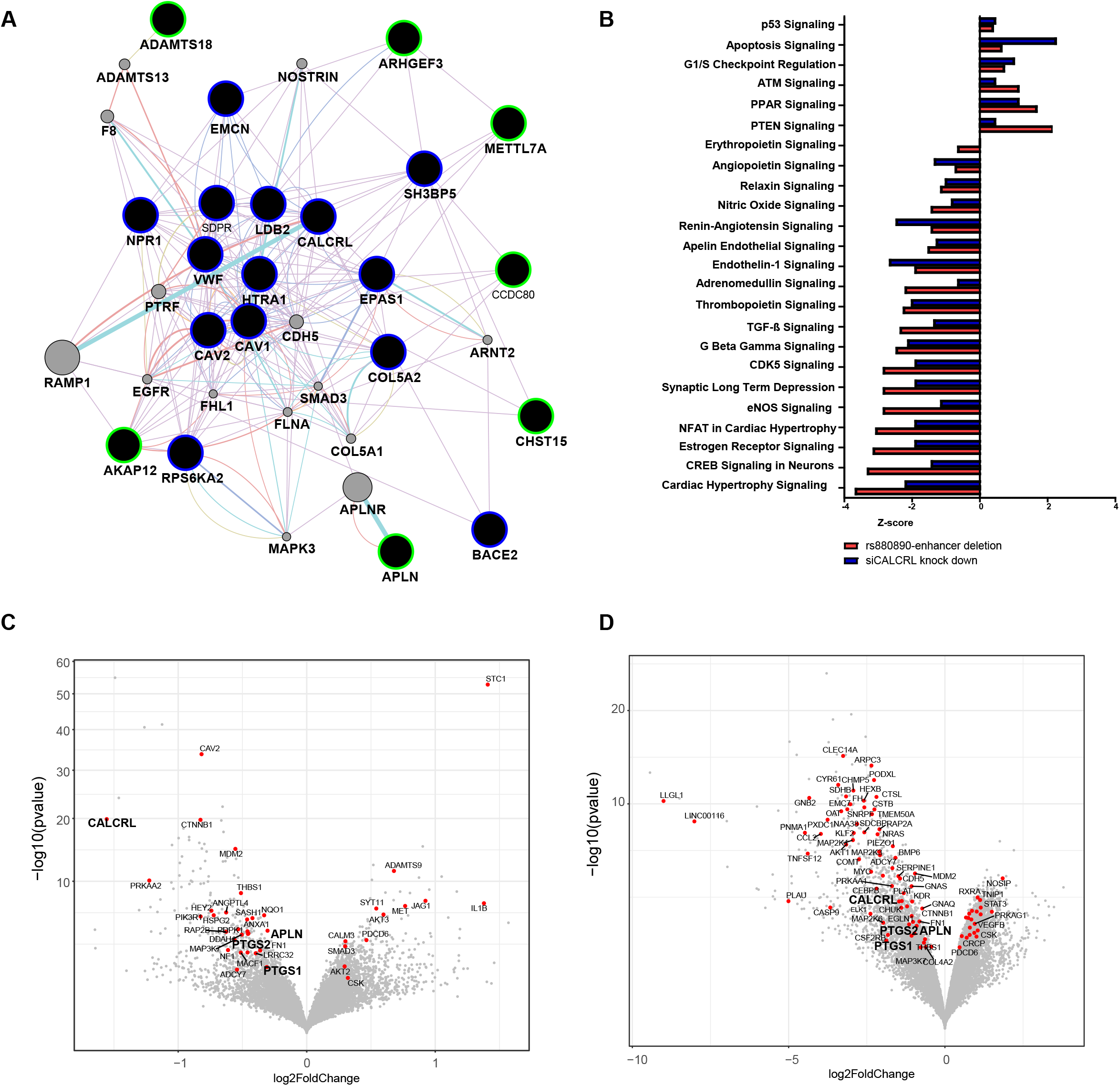
**A)** Network generated by Cytoscape app geneMANIA. Black nodes with blue outline represent known *CALCRL* co-expressed genes while black nodes with green border are novel co-expressed genes. Purple connections denotes co-expression, cyan denotes pathway, pink denotes physical interaction, purple denotes co-localization. **B)** Functional enrichment plot of *CALCRL* siRNA and rs880890-enhancer deleted samples using IPA. z-score indicates a predicted activation or inhibition of a pathway (i.e. Erythropoietin, Angiopoietin, Relaxin, Nitric Oxide, Renin-Angiotensin, Apelin Endothelial, Endothelin-1, Adrenomedullin, Thrombopoietin and eNOS signaling pathways are predicted to be repressed). **C-D)** Volcano plot highlighting selected candidates differentially expressed in RNA-Seq data from *CALCRL* siRNA and rs880890-enhancer deleted samples.

**Figure 5.**
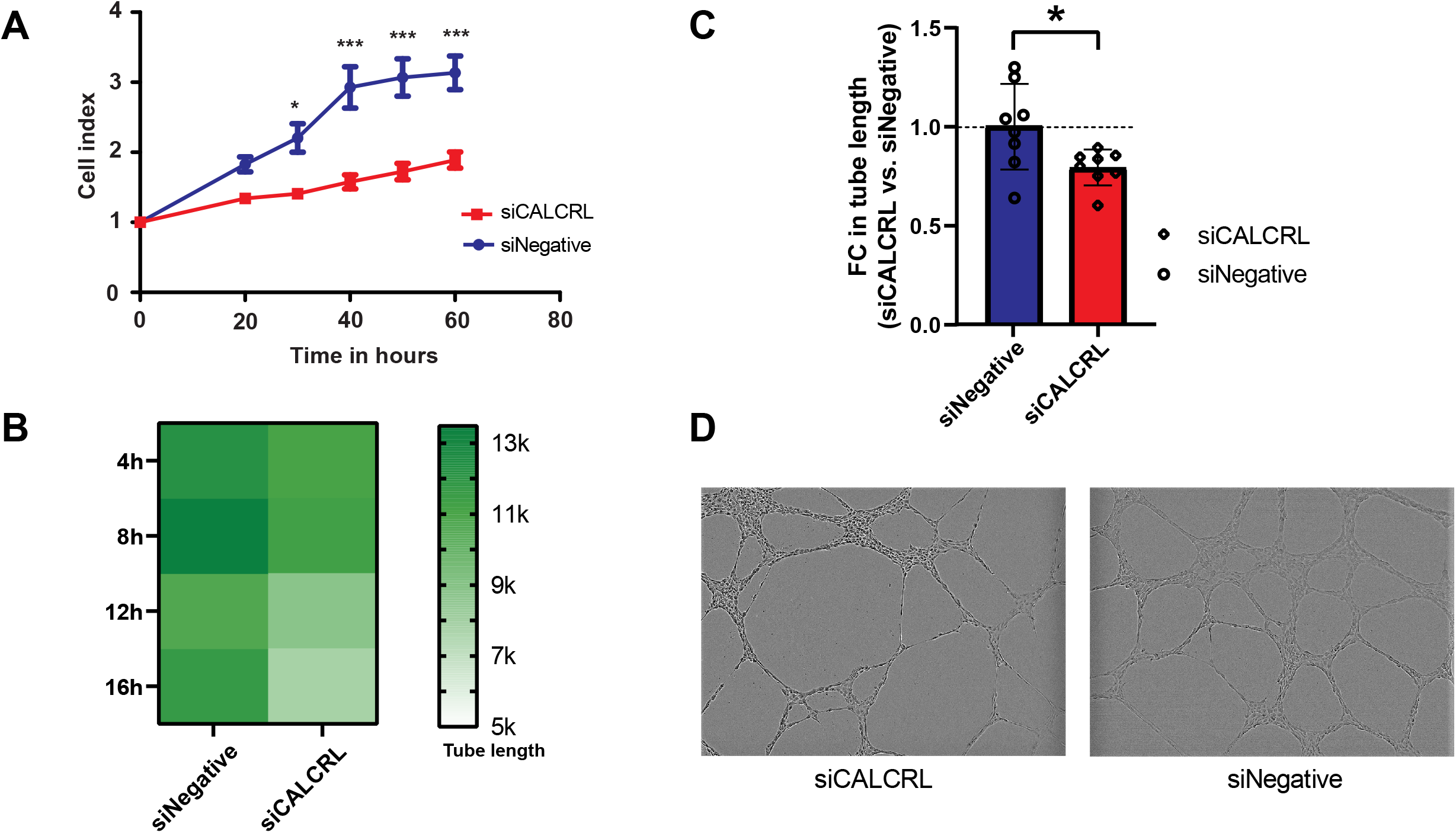
**A)** Effect of *CALCRL* downregulation on proliferation of TeloHAECs. Normalized cell index (xCELLigence system) upon siRNA knockdown of *CALCRL* in TeloHAECs shows significant decrease in cell proliferation. **B)** Effect of CALCRL knockdown on tube formation. Heat map of the averaged total tube lengths 4–16 h after plating of TeloHAECs on Matrigel (n=3). **C)** Fold change in tube length formation. Each bar represents the average + SEM of total tube lengths obtained from each-well image. Two-way ANOVA with of *CALCRL* siRNA treatment vs scrambled negative controls at each time point: *P < 0.05; **P < 0.01; ***P < 0.001; Data points are average values of three biological replicates. Tube length values comparing siCALCRL to siNegative was used analyzed by 2-tailed Student t test. *P<0.05. **D)** Representative of 9 spot well output images from Incucyte after 24 h siCALCRL treatment (n=3).

**Figure 6.**
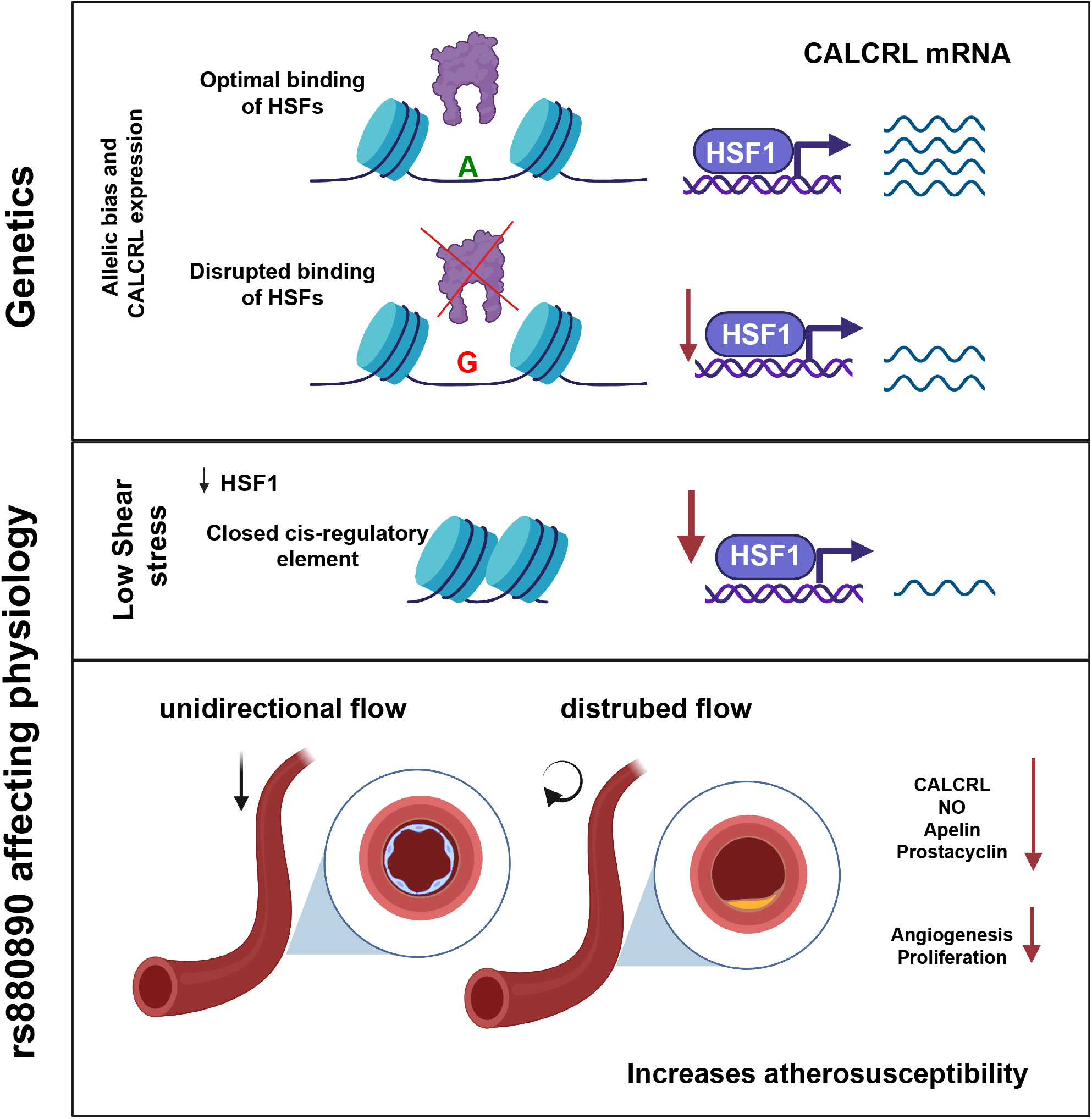
Schematic model depicting the interplay between genetics and hemodynamic forces at *CALCRL* loci and rs880890 at the molecular level in regulating endothelial *CALCRL* expression and vascular functions.

Further analysis of the differentially expressed genes (DEGs) identified 390 and 990 genes significantly regulated (padj<0.05) upon gene silencing or enhancer deletion, respectively (**Table S9**). Importantly, this included 25% of the 195 *CALCRL* co-expressed genes, demonstrating significant enrichment (hypergeometric t-test P-value=6.5E-07). Functional enrichment analysis demonstrated that similar pathways were regulated by both the gene and enhancer repression (**Figure 4B**). The most important pathways affected included the repression of apelin, endothelin, adrenomedullin, eNOS and angiopoietin signaling pathways. Apelin (*APLN*), Cyclooxygenase (*PTGS1* and *PTGS1*) genes themselves were significantly downregulated in both RNA-Seq experiments indicating they could be direct downstream targets of CALCRL **(Figure 4C-D)**. Among DEGs genes, *PTGS1* and *PTGS2* were also downregulated in disturbed flow compared to unidirectional flow further linking these genes to shear stress (**Suppl. Figure 7**). Interestingly, we also observe a downregulation trend of *EDN1*. The RNA-Seq results also found that depletion of *CALCRL* expression leads to repression of TGF-β signaling (*THBS1, PTGS2, and TGFBR2*) and angiogenesis regulatory genes (*PDPK1, ANGPTL4*). In line with this, knock down of *CALCRL* led to a significant decrease in cell proliferation (**Figure 5A**) and tube formation (**Figure 5B through D**).

## Discussion

Blood flows through the arteries with tangential force that acts on the surface of endothelial cells causing shear stress^56^ Shear stress spans a range of spatiotemporal scales and contributes to regional and focal heterogeneity of endothelial gene expression, which is important in vascular pathology^57^. To this end, exposure to disturbed blood flow at arterial bifurcations and curves causes constitutive activation of vascular endothelium contributing to atherosclerosis, the major cause of coronary artery disease (CAD)^58^. It has been known that endothelial cells undergoing shear stress develop atherosclerotic plaques due to disruption in eNOS pathway. Krause *et al*.^59^ recently reported that the noncoding common variant at rs17114036, associated with CAD/IS in GWAS, regulates *PLPP3* expression in endothelium through increased enhancer activity that is dynamically regulated by unidirectional flow and transcription factor KLF2. Here, we extend the discovery and characterization of genetic variants associated with CAD acting through shear stress regulation, by identifying a mechanosensitive endothelial enhancer that regulates *CALCRL* expression. Specifically, we identified a non-coding, common genetic variant rs880890 that modified levels of *CALCRL* expression in endothelial cells. We demonstrate that rs880890 confers increased enhancer activity that is dynamically regulated by unidirectional flow and HSF transcription factor(s).

Our data suggests that decreased endothelial production of CALCRL could promote atherosclerosis. This directionality of effect has been previously experimentally validated in mice, where endothelial-specific deficiency of CALCRL lead to increased atherosclerotic lesions^56^. In addition to CAD, 2q32.1 locus is also associated with hypertension. In line with this, the G allele at rs880890 is associated with increased risk for hypertension^60^. Since G allele of rs880890 reduces CALCRL production, that is normally needed increase in cAMP levels, activate protein kinase A (PKA) and eNOS^13^, deficient NO formation could explain its association with hypertension. Interestingly, the minor allele frequency (G) of rs880890 is highly variable between the ethnicities ranging from 0.2 in the African population to 0.9 in Asian populations such as Japanese^61^ suggesting population specific differences in the risk susceptibility mediated by this locus.

Vasodilation is one of the important characteristics of blood vessels to aid in reducing the stress put on the borders of the vessel by the blood flow. eNOS derived NO is an endogenous vasodilatory gas that continually regulates the diameter of blood vessels and maintains an anti-proliferative and anti-apoptotic environment in the vessel wall^62^. Our data from perturbation experiments demonstrate that CALCRL could play a major role in regulation of several vasodilatory factors. To this end, we identify apelin (APLN)^63^, endothelin (EDN1)^64,65^ as potential downstream target of CALCRL. In addition, PTGS1 (COX-1) and *PTGS2* (COX-2), key enzymes which convert arachidonic acid to prostaglandins to mediate vasodilation and inhibition of platelet aggregation,^66^ were identified. Further investigations are warranted to shed light on the involvement of CALCRL in regulating the expression of *APLN, PTGS1, PTGS2* and *EDN1*.

CALCRL/adrenomedullin/G-protein-alpha activation in endothelial cells have been shown as a promising approach to inhibit progression of atherosclerosis by reducing endothelial inflammation^56^. In this study, we provide evidence that *CALCRL* expression is regulated by the concerted action of genetic variation and shear stress to promote pathogenic mechanisms leading to plaque formation (**Figure 6**). We present a model where the ‘G’ allele of the rs880890 associated with increased risk of CAD confers decreased endothelial enhancer activity thereby reducing *CALCRL* expression. Under disturbed flow, Changes in histone acetylation limits DNA accessibility, further reducing HSF1/2/4 binding and *CALCRL* expression. Changes in *CALCRL* expression may influence vasoconstriction and atherosclerotic plaque formation through the regulation of eNOS, apelin, adrenomedullin, renin-angiotensin, angiopoietin and endothelin-1 signaling pathway with reduced expression promoting pathology. In summary, this study has identified a previously unreported mechanosensitive pathway, and HSFs as important transcription factors that exert an anti-atherosclerotic effect in endothelial cells. Our data highlights the utility of using (GWAS) data to illuminate the molecular mechanisms that drive pathology and provides a plausible mechanism for how human SNPs in *CALCRL* regulates cardiovascular disease.

## Data availability

The RNA-Seq experiments reported in this study are deposited in the GEO database under the accession number: GSE222118 and will be made public upon manuscript acceptance. Reviewers can access the submission using the following secure token: mjwrimggplanpif.

## Acknowledgements

The authors wish to acknowledge Biocenter Finland for infrastructure support, CSC – IT Center for Science, Finland and Bioinformatics center of University of Eastern Finland for the computational resources.

This research was supported by the European Research Council (ERC) under the European Union’s Horizon 2020 research and innovation program (Grant No. 802825 to M.U.K), the Academy of Finland (Grants Nos. 287478 and 319324 to M.U.K.), the National Institutes of Health (NIH) (R01HL147187/HL/NHLBI NIH HHS/United States to C.E.R.)., American Heart Association (20PRE35200195 to L.K.S.). Academy of Finland (276634 and 312487 to M.H and P.P), Instrumentarium Science Foundation to I.S, the Finnish Foundation for Cardiovascular Research, the Sigrid Juselius Foundation, and the Doctoral Program of Molecular Medicine at University of Eastern Finland.

---

**Suppl. Figure 1:**
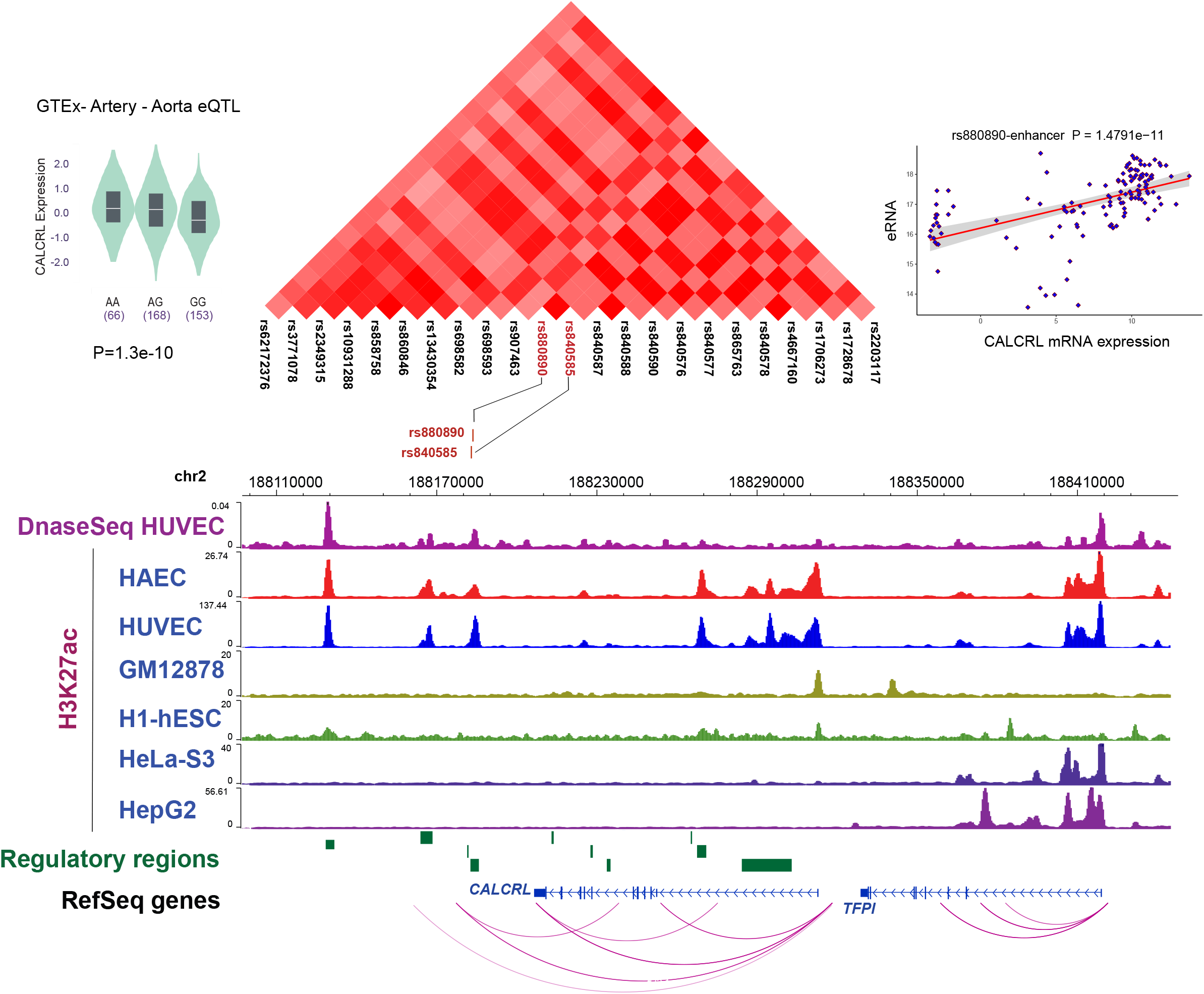
Effects of regulatory SNP (rs880890) and target gene (CALCRL) across multiple data in endothelial cells. A) The CALCRL eQTL at rs880890 is shown in Aorta from GTEx. B) Washu browser shot of CAD loci with CALCRL region showing Promoter capture Hi-C interactions between promoter of CALCRL and regulatory regions. Analysis of H3K27ac signal across several cell types demonstrating the selection of endothelial specific active regulatory elements included in this study. C) Plot showing the correlation of GRO-Seq eRNA and CALCRL mRNA expression. A positive correlation with a P-value of 1.38E−71 is observed

**Supplementary figure 2:**
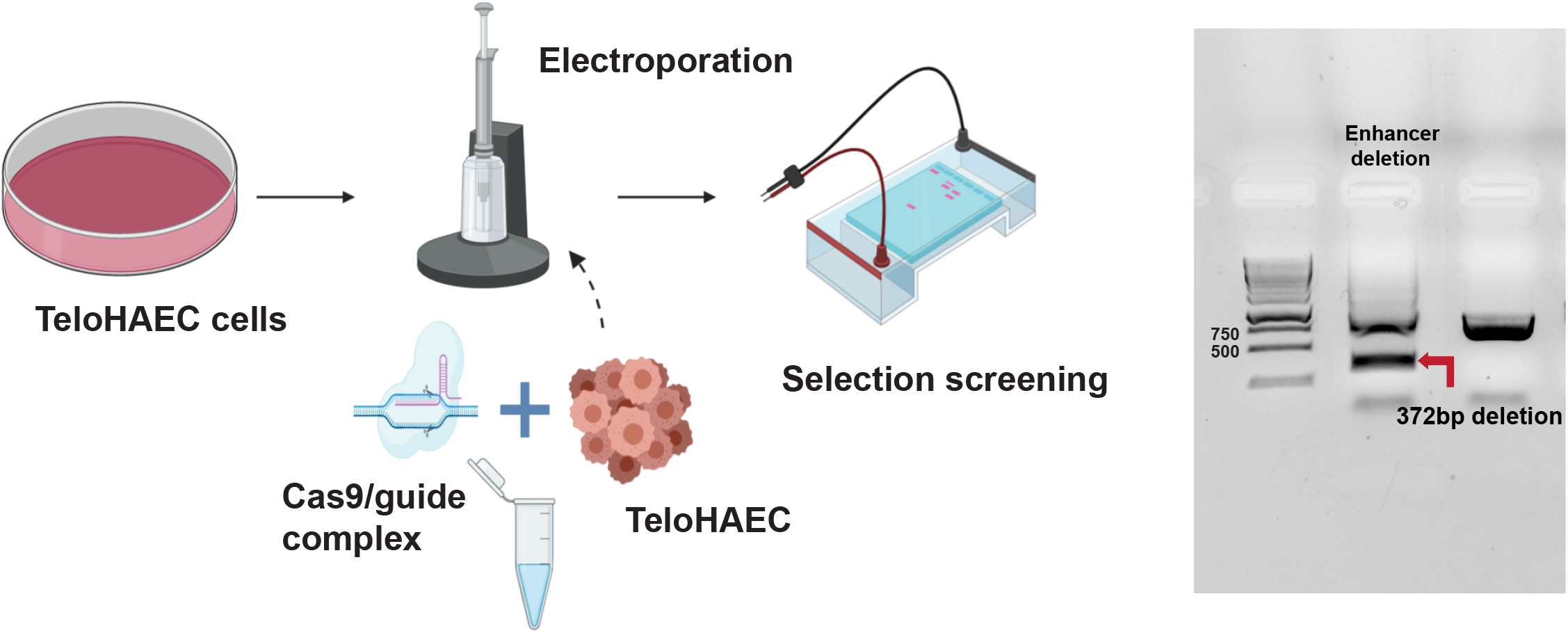
CRISPR-mediated deletion of the risk variant carrying enhancer in TeloHAEC cells. A) Schematic representation of CRISPR-Cas9 enhancer deletion experiment. B) PCR analysis of the deletion efficiency showing the deletion of 372bp of the rs880890-harboring enhancer in TeloHAECs.

**Suppl. Figure 3:**
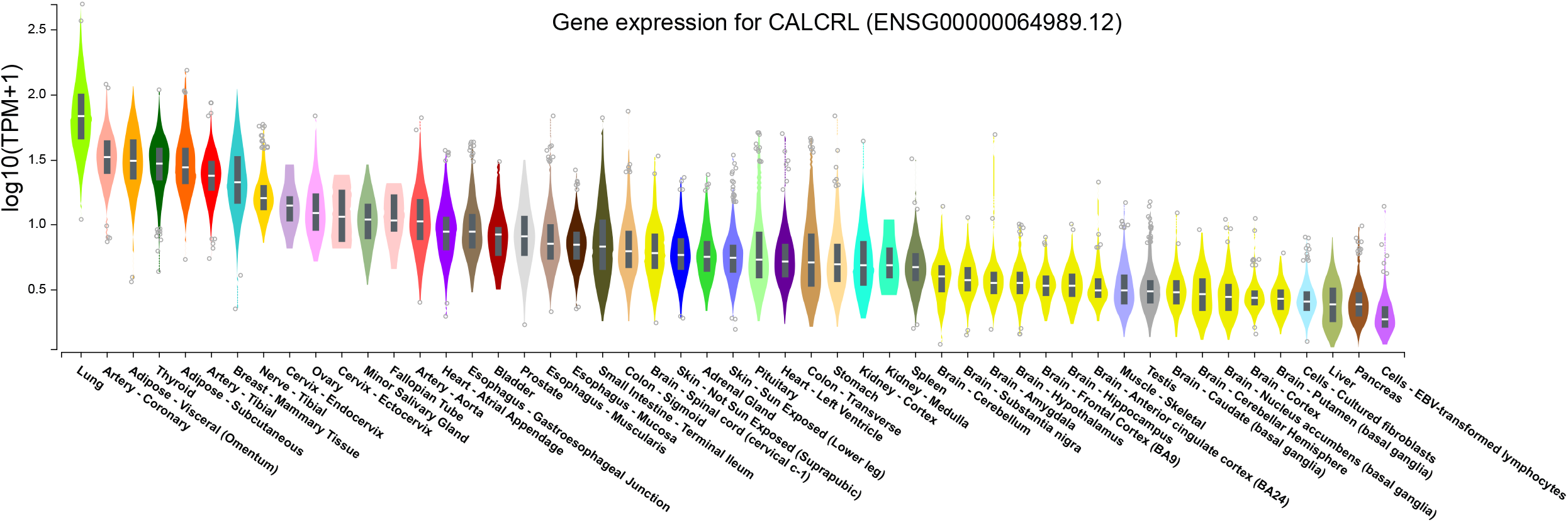
Gene expression of CALCRL across multiple human tissues form GTEx release V8. Expression values are shown in TPM (Transcripts per million), calculated from a gene model with isoforms collapsed to a single gene. Box plots are shown as median and 25th and 75th percentiles; points are displayed as outliers if they are above or below 1.5 times the interquartile range. The values are shown in logarithmic scale sorted by their median values.

**Suppl. Figure 4:**
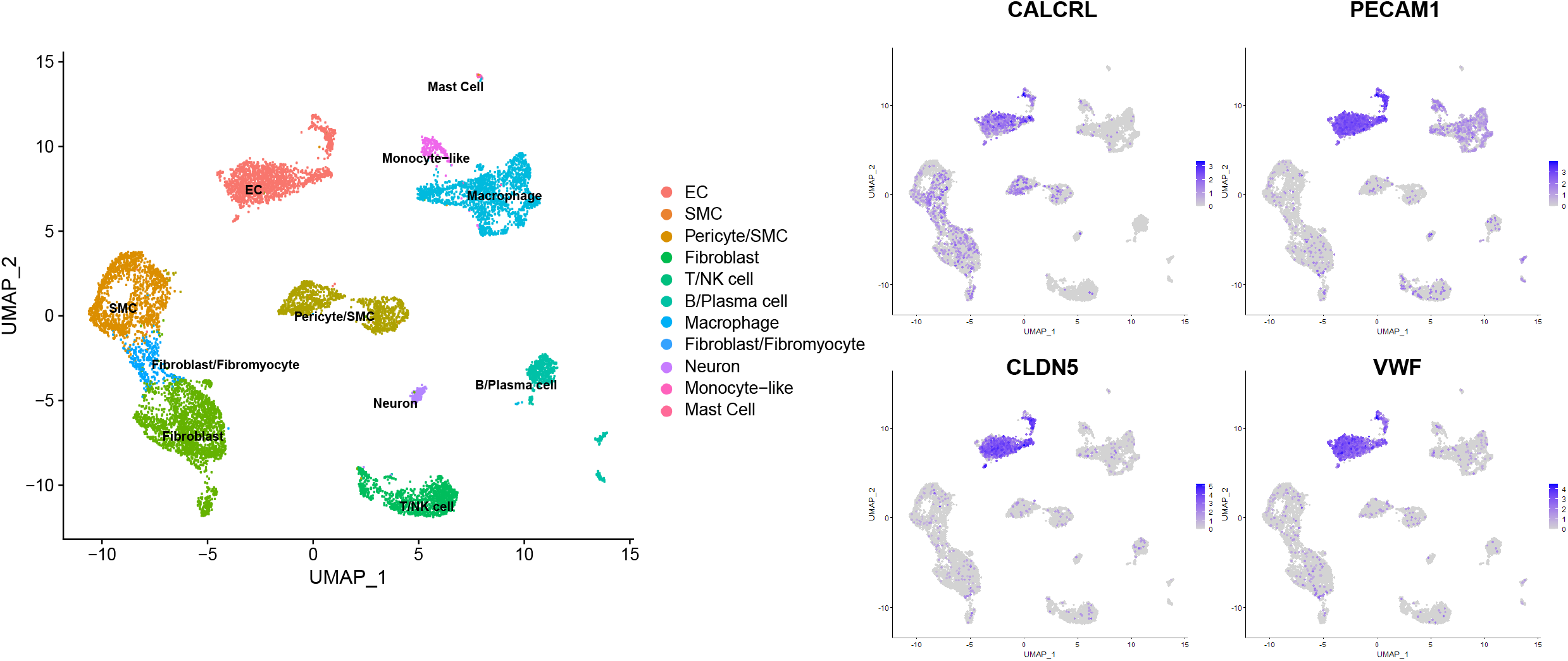
Single cell RNA-Seq data from human coronary arteries showing the different cell type clusters.^21^ We can see that along with other endothelial specific markers, CALCRL is predominantly expressed in endothelial cells.

**Suppl. Figure 5:**
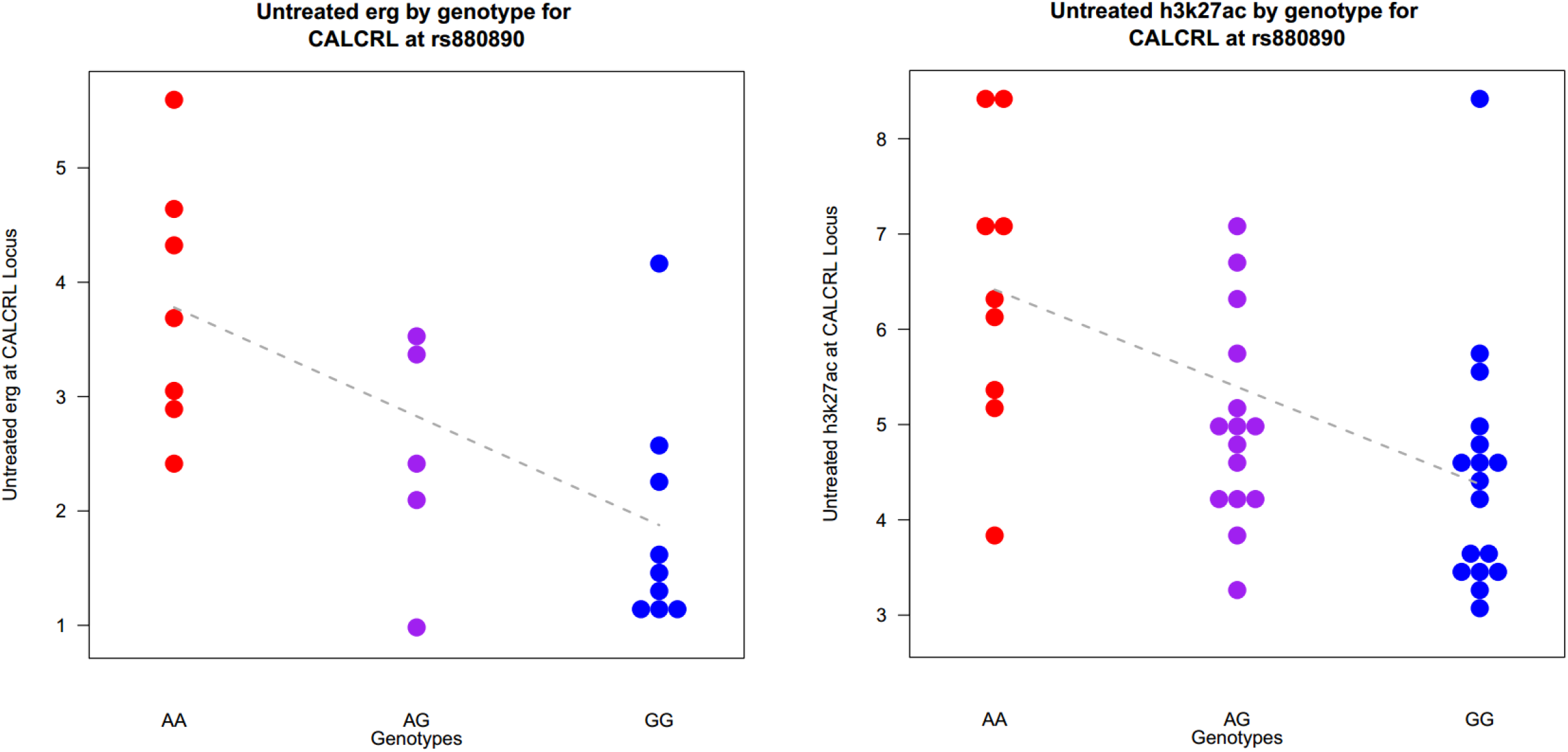
A trend of allele specific deposition of histone mark H3K27ac and the binding of endothelial specific transcription factor ERG was also detected in a cohort of HAEC donors

**Suppl. Figure 6:**
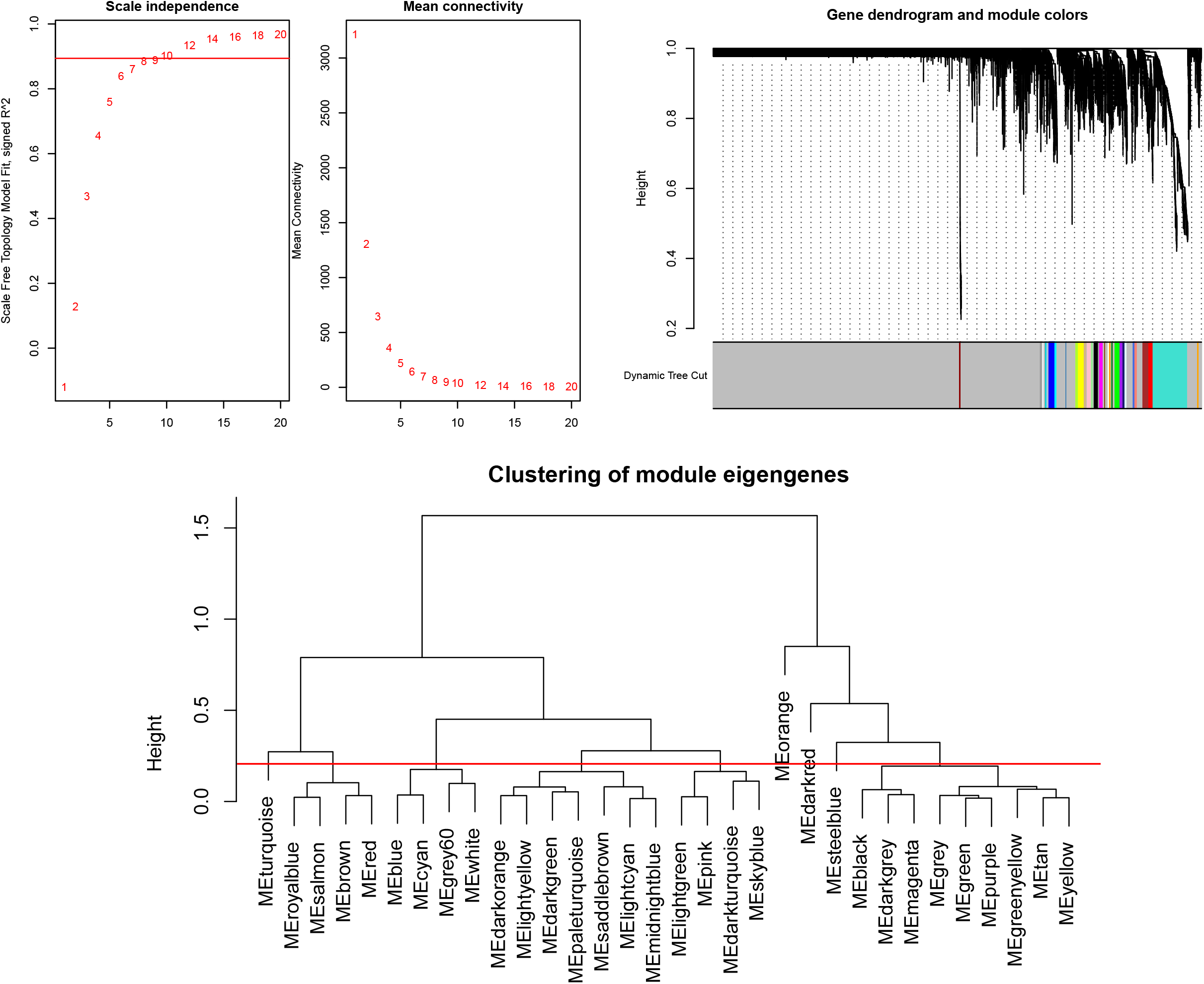
A) Analysis of network topology for various soft-thresholding powers. The left panel shows the scale-free fit index as a function of the soft-thresholding power (x-axis). The right panel displays the mean connectivity as a function of the soft-thresholding power (x-axis). B) Clustering dendrogram of genes, with dissimilarity based on topological. C) Clustering dendrogram of genes, with dissimilarity based on topological overlap. Modules whose expression profiles are very similar were merged with a height cut of 0.1.

**Suppl. Figure 7:**
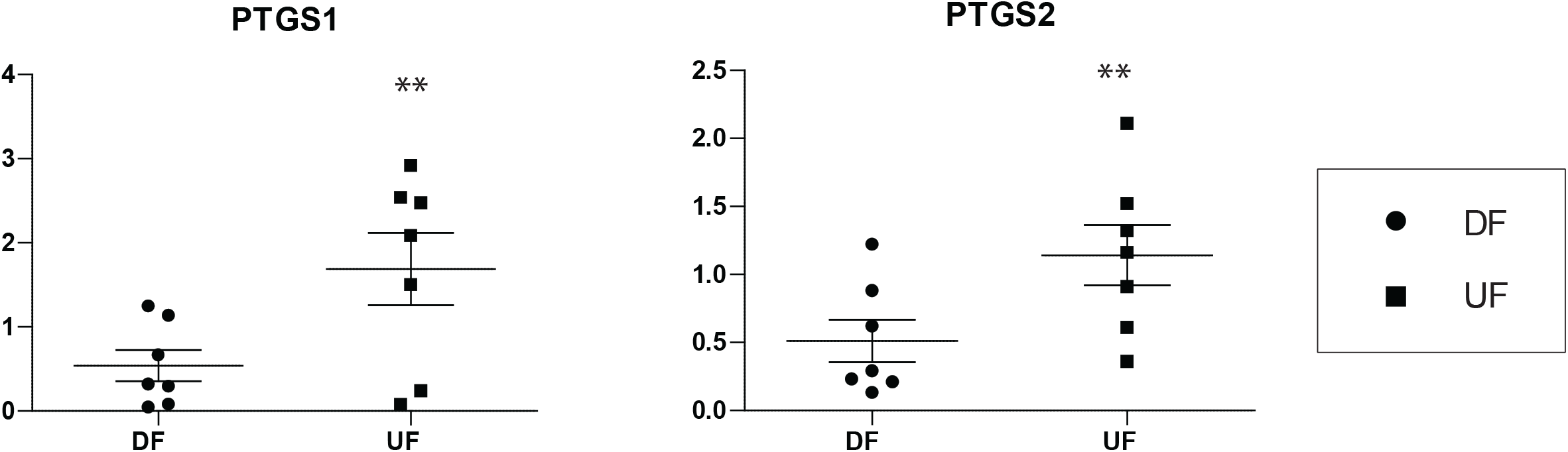
qPCR on mRNA expression of PTGS1 (P=0.0073) and PTGS2 (P=0.0072) under shear stress conditions seven donors of human coronary artery endothelial cells. Both genes were significantly downregulated in disturbed flow compared to unidirectional flow.

## Notes

### Competing Interest Statement

The authors have declared no competing interest.

